# Specific histone modifications associate with alternative exon selection during mammalian development

**DOI:** 10.1101/361816

**Authors:** Q Hu, CS Greene, EA Heller

## Abstract

Alternative splicing (AS) is frequent during early mouse embryonic development. Specific histone post-translational modifications (hPTMs) have been shown to regulate exon splicing by either directly recruiting splice machinery or indirectly modulating transcriptional elongation. In this study, we hypothesized that hPTMs regulate expression of alternatively spliced genes for specific processes during differentiation. To address this notion, we applied an innovative machine learning approach to relate global hPTM enrichment to AS regulation during mammalian tissue development. We found that specific histone modifications, H3K36me3 and H3K4me1, play a dominant role in skipped exon selection among all the tissues and developmental time points examined. In addition, we used iterative random forest model to identify interactions of several hPTMs that associated with skipped exon selection during tissue development. Collectively, our data demonstrated a link between hPTMs and alternative splicing which will drive further experimental studies on the functional relevance of these modifications to alternative splicing.

## Background

Alternative splicing (AS) is a regulatory mechanism of gene expression that enables one gene to generate multiple mRNA isoforms that may have different functions or properties. RNA-seq analyses of the whole transcriptome have revealed the high prevalence of AS in many organisms (human and mouse: 90%, drosophila: 60%) [1, 2]. AS has been shown to contribute to cell differentiation, tissue identity and organ development [2]. The expression of a specific isoform is often necessary to maintain tissue identity and function, while selection between alternative isoforms drives tissue development and cell differentiation [3]. Understanding the regulation of developmental processes requires the investigation of AS events across different tissues during development. A number of studies aimed at revealing the importance of AS during development find that AS and specific isoform expression is frequent during early mouse embryonic development [4-6]. In addition, many alternatively spliced isoforms show a dramatic change in relative expression levels during embryonic to adult development in *C.elegans* [7]. Studies targeted at the underlying mechanism of AS regulation have largely identified which splice motifs interact with the splicing machinery to facilitate and regulate splicing. Several AS regulators that are critical to tissue development have been identified, such as CELF1 in heart development [8], ELAVL, PTBP1 and NOVA1/2 in brain development [9-11] and ESRP1 in liver development [12]. However, as these elements are not sufficient to explain all aspects of AS regulation, including specific gene targeting, additional regulatory mechanisms must exist to direct the selection of alternatively spliced isoforms [13].

In addition to specific gene sequences that specify transcriptional and splicing motifs, epigenetic mechanisms function in transcriptional regulation and play important roles in many biological processes [14, 15]. Genome regulatory elements undergo dynamic changes in the enrichment of histone post-translational modifications (hPTMs), which function during development to direct expression of corresponding genes [16, 17]. For example, co-enrichment of H3K4me3 and H3K27ac is found at enhancers related to heart development in mouse [18] and functions to regulate the expression patterns of genes involved in developmental transitions in the cardiac lineage [19]. In addition, computational analysis in several cell lines has found that particular hTPMs enriched around the transcriptional start sites of expressed genes associate with transcription initiation [20, 21], while the levels of H3K4me3 and H3K79me1 are associated with steady-state expression of particular exons and genes [21-23].

In addition to the above roles of hPTMs in regulating gene expression, recent evidence suggests that hPTMs also function in the specification of exons spliced into a transcribed gene [24, 25]. Specific hPTMs have been proven to regulate exon splicing by either directly recruiting splicing factors and adapters or indirectly modulating the elongation rate of RNA polymerase II (RNAPII), indicating a potential link between histone modification and alternative splicing [25, 26]. Studies on human datasets show that several types of hPTMs are associated with exon inclusion or exclusion. A recent study in human stem cells shows specific hPTMs, such as H3K36me3, regulate alternative splicing events and are involved in nonsense-mediated mRNA decay of BARD1 (BRCA1-associated RING domain protein 1). The authors also compare the contribution of genomic features and epigenetic features to alternative splicing and find that epigenetic features are more important to differentiate different splicing patterns [27].

Due to the critical role of AS in tissue development and the potential link between specific hPTMs and AS in ES cell differentiation, we hypothesized that hPTMs could also drive development by regulating expression of alternative spliced genes for specific processes in mammalian tissue development. To address this notion, we utilized a state-of-the-art machine learning approach to conduct a genome-wide, analysis that relates hPTMs to AS regulation during mammalian tissue development. We integrated ChIP-seq and RNA-seq data from 7 different mouse embryonic tissues at 6 developmental time points to determine (1) which hPTMs associate with alternatively spliced exons, (2) which hPTM(s) play dominant role(s) in alternative exon selection and (3) the interaction of multiple hPTMs in exon selection. We focused on one specific alternative splicing type – skipped exon – because it is the most prevalent alternative splicing event in mammalian tissue and contributes greatly to proteome diversity [28]. We categorized two subtypes of skipped exons based on RNA-seq data analysis: developmental gain/loss and isoform selected high/low. Skipped exons were categorized as developmental gain/loss if the isoform switch occurred during development. Alternatively, skipped exons were categorized as isoform selected high/low if their inclusion was in the upper (75%) and lower (25%) quantiles, respectively, and isoform expression did not change over development. Enrichment analysis found these two groups of AS genes consisted of different functional categories. We also observed that the number of AS events increased over developmental time, with brain tissue showing the most significant increase. To infer the relevance of hPTMs to AS events across tissues and development, we analyzed the ChIP-seq signal distribution of 8 distinct hPTMs (H3K36me3, H3K4me1, H3K4me2, H3K4me3, H3K27ac, H2K27me3, H3K9me3 and H3K9ac) in the exon-flanking region. Remarkably, we found that only two hPTMs, H3K36me3 and H3K4me1, were differentially enriched with respect to skipped exon category.

We further derived a computational model for predicting skipped exon category using hPTM signal in the exon flanking regions. We found that hPTMs can accurately predict skipped exon category in both developmental gain/loss and isoform selected high/low groups, indicating the potential link between hPTM and skipped exon selection. Our finding indicates that specific histone modifications, H3K36me3 and H3K4me1, play a dominant role in skipped exon selection among all the tissues and developmental time points examined. Furthermore, the contribution of some hPTMs was tissue-specific. In brain tissues and heart, H3K9ac had a relatively higher predictive rank, while in limb, neural tube and liver, the effect of H3K27me3 was higher. We also identified interactions of two or more hPTMs that highly predict AS. For example, the interaction between H3K36me3 and H3K4me1 in the exon flanking region was the top feature in both skipped exon categories. The other top interactions included H3K27me3/H3K36me3, H3K27ac/H3K36me3, H3K27ac/H3K4me1, and H3K36me3/H3K9me3. Collectively, our data demonstrate a link between hPTM and alternative splicing in mouse tissue development, which will drive further experimental studies on the functional relevance of these modifications to alternative splicing.

## Results

### Characterization of alternative spliced events in tissue development

Alternative splicing has been shown to contribute to cell differentiation, tissue identity and organ development [2, 29]. To identify AS events associated with tissue development, we analyzed ENCODE RNA-seq data [30] derived from mouse embryonic tissues at multiple developmental time points. We selected data from 7 tissues at 6 time points based on the availability of both RNA- and ChIP-seq data. Our analysis focused on one specific AS type – skipped exon – because it is the most common type in the mammalian transcriptome [28]. Significant skipped exon events in each dataset were identified by comparing each time point with the earliest time point (E11.5) using rMATS [31]. We analyzed alternative splice events over developmental time for each tissue and identified skipped exons with significant ΔPSI larger than 0.1 (developmental gain) or less than −0.1 (developmental loss) (FDR < 0.05). These skipped exons associated with tissue development are referred to as ‘developmental gain/loss,’ and vary in number from 600 to 3000 across the tissues examined (Figure 1A, Supplemental Table 2).

**Figure 1.**
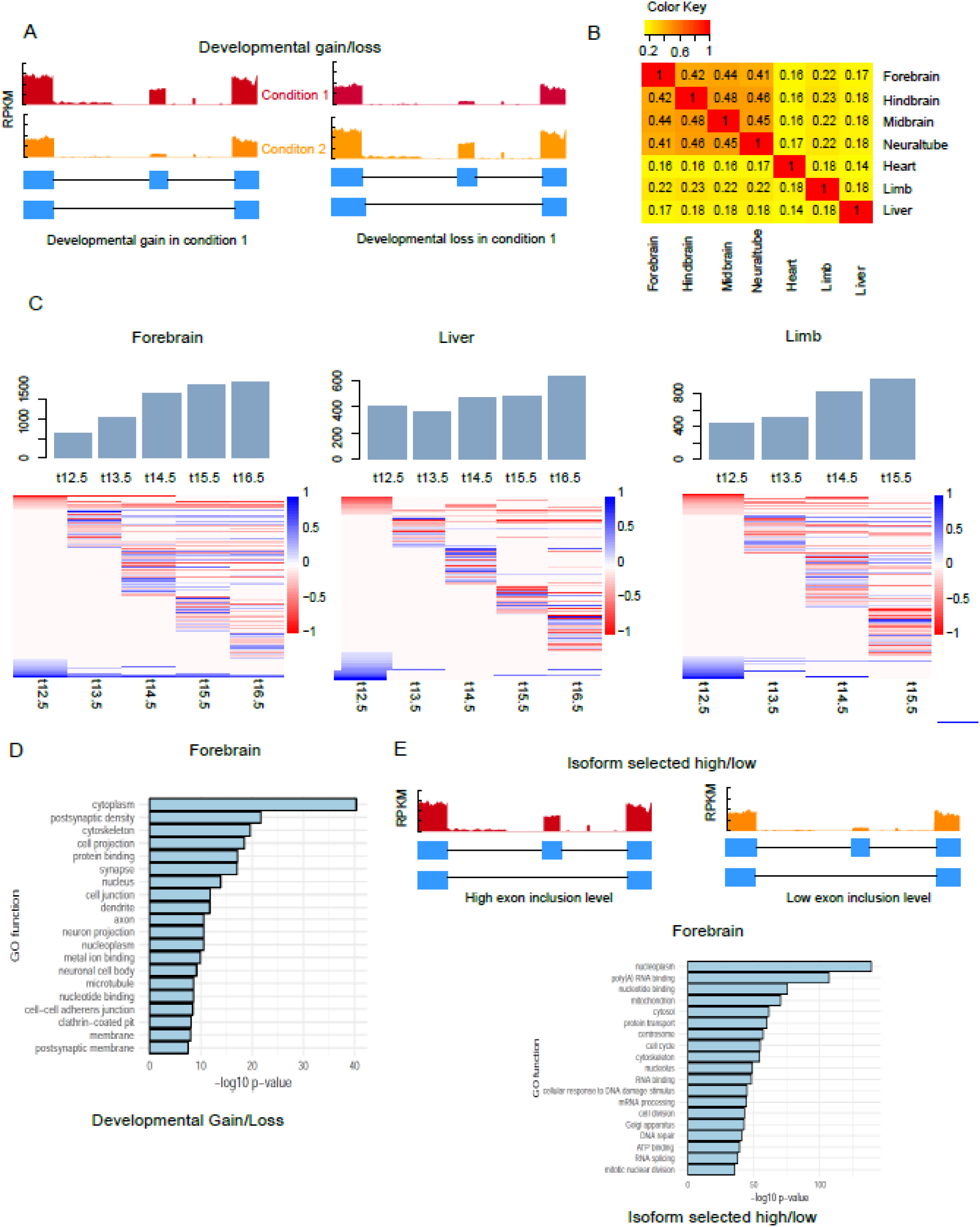
Alternative splicing events associated with tissue development. (A) Schematic illustrating developmentally associated skipped exons. (B) Overlap of developmentally associated alternative splicing events between tissues (C) Number of significant alternative splicing events identified across 7 developmental time points for several mouse tissues. Heat maps show the differential exon inclusion level (ΔPSI) by comparing each time point to earliest time point in skipped exons (row). Bar graph shows the proportion of lineage-specific AS genes. (D) Ontology analysis of developmentally associated AS genes in forebrain. Data are derived from ENCODE database [30] (supplemental table 1) and analyzed using rMATs [31].

We also observed that the number of developmentally associated alternative splicing events increased with developmental time, with brain tissues showing the greatest increase (Figure 1C, Supplemental Table 2, Supplemental Figure 1). For example, the number of alternative splicing events in forebrain increased 65%, from 625 on E12.5 day to 1941 on E16.5, while in liver the increase was only 36% (405 to 637). Hierarchical clustering on skipped exon events across developmental time points revealed specific splicing patterns in different tissues (Figure 1C, Supplemental Figure 1). In brain, neural tube and limb, there were more developmental gain events after E12.5, while in heart and liver, the number of developmental loss events was slightly higher at most time points (Supplemental Table 2). Tissue-specific alternative splicing plays important roles for tissue identity during development [32]. Thus, to explore what percentage of developmentally-associated skipped exons are tissue-specific, we performed pairwise comparison of the identified skipped exons among different tissues (Figure 1B). On average, over half of the lineage-specific transcripts in each tissue were alternatively spliced; this percentage was not significantly different between tissues (Supplemental Figure 1). In addition, we found most of the lineage-specific events occurred in the early time point, which account for ~50% of those events among all tissues. Brain tissues showed a significant decrease of lineage-specific events at later time points when compared to liver, limb and heart (Supplemental Figure 1). This finding underscores the relevance of alternative splicing to lineage-specific gene expression.

In addition to developmentally associated skipped exons, we also observed another category of skipped exons according to PSI values derived from rMATS – isoform selected high/low (Figure 1E). These exons were alternative spliced but did not show inclusion level changes over developmental time.

Gene ontology (GO) enrichment analysis found that these two categories of AS genes were enriched in different functional categories (Figure 1D, 1E, Supplemental Table 3). AS genes belonging to the developmental gain/loss category were overrepresented in certain GO functions, such as cytoplasm, postsynaptic density and cytoskeleton (Figure 1D), consistent with previous studies of AS genes in mouse tissue development [33, 34]. Alternatively, AS genes belonging to the inclusion high vs low category were enriched in most of the other functional categories, such as RNA binding, cell cycle and cell division (Figure 1E). Taken together, these results comprised a global analysis of alternative splicing events in different tissues across development.

### Histone modification enrichment in exon flanking regions differentiated skipped exon groups

Though previous studies find that histone modifications are enriched in promoter regions and predict expression of corresponding genes [21, 23, 35], it has become increasingly clear that they also associate with gene bodies and exon regions, indicating a potential role of histone modifications in pre-mRNA splicing regulation [36]. To investigate if histone modifications associated with alternative splicing across tissue development, we focused on the ChIP-seq distribution patterns of 8 histone modifications, including H3K4me1, 2, 3, H3K9me3, H3K27me3, H3K36me3, H3K9ac and H3K27ac, which were available for all tissues and developmental time points analyzed. For each developmental time point, we profiled the hPTM distribution patterns of all skipped exons. We reasoned that histone modifications related to alternative splicing are likely to be localized to the genomic region at which splicing occurs and hypothesized that ChIP-seq distribution patterns would vary by skipped exon category. Thus, we compared the normalized ChIP-seq signal distributions of each hPTM in a +/- 150 bp region flanking the splice sites of each skipped exon.

Figure 2 shows the mean ChIP-seq signal distributions of several hPTMs in brain and heart (the distributions of all 8 hPTMs in all 7 tissues at each developmental time point are in Supplemental Figures 3-36). We found that only two modifications, H3K36me3 and H3K4me, distributed according to skipped exon category. In addition, hPTMs corresponding to different groups of skipped exons diverged greatly with respect to their correlative behavior. For example, H3K36me3 was positively correlated with exon inclusion levels of skipped exons in the isoform selected high vs low inclusion category. That is, the higher the exon inclusion level, the stronger the H3K36me3 enrichment in the exon flanking regions. However, H3K4me2/3 displayed the opposite trend, that is, the H3K4me2/3 enrichment was highest in skipped exons with low inclusion level. This association pattern was consistent among all tissues (Figure 2, Supplemental Figure 3-36). Conversely, for the gain vs loss inclusion category, the association patterns were similar in brain tissues but differed among the other tissues. This is especially true for H3K4me2/3, as we found that in forebrain at E12.5, H3K4me2/3 enrichment was positively correlated with skipped exons with inclusion gain, but in heart, limb and liver, enrichment appeared to be negatively correlated with those exons.

**Figure 2.**
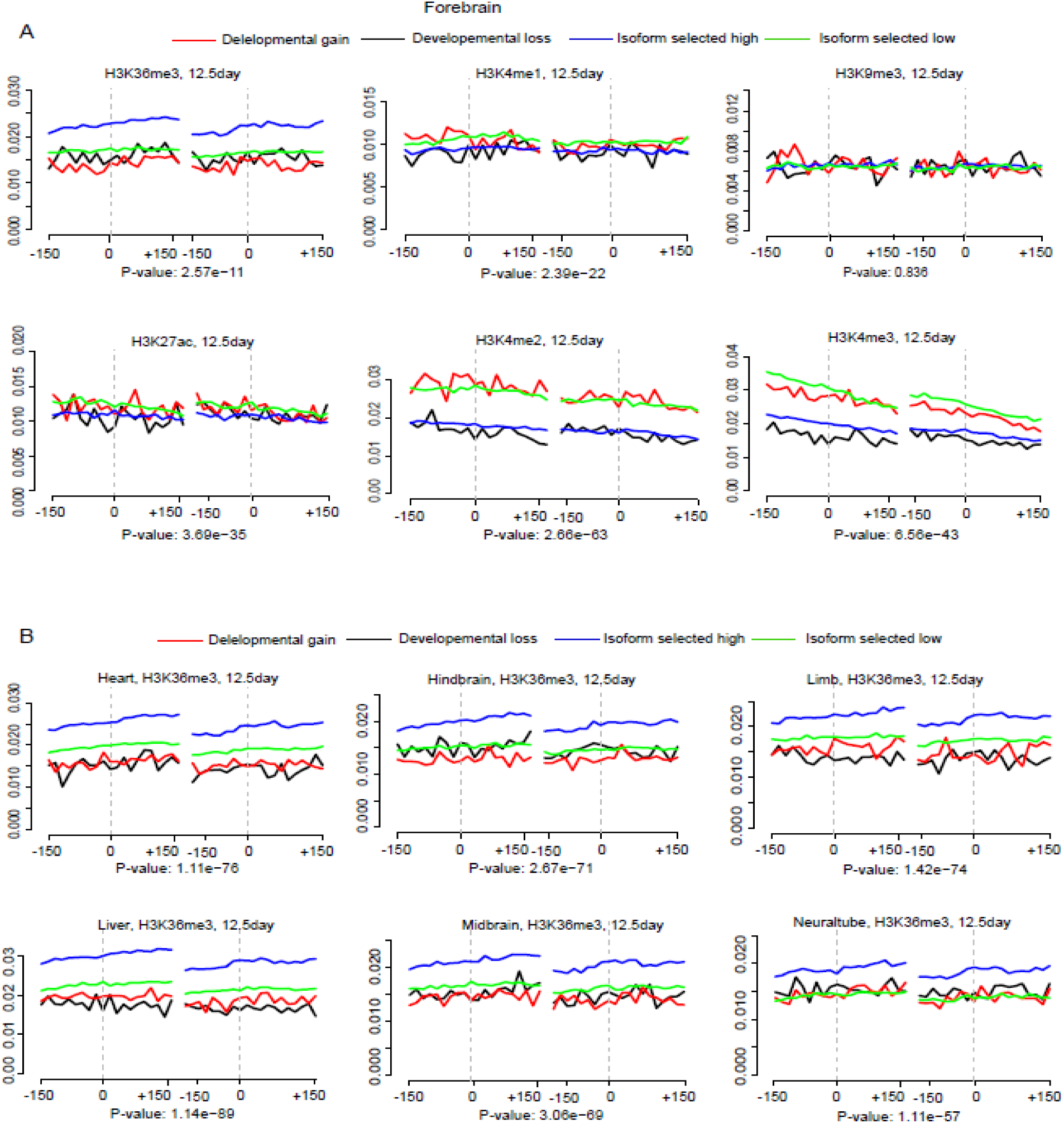
Representative distribution of mean ChIP-seq signal of 6 types of hPTM, including H3K36me3, H3K4me1, H3K9me3, H3K27ac, H3K4me2 and H3K4me3 on the flanking region (+/- 150bp) of four types of skipped exons. Dashed grey line shows exon-intron borders. (A) forebrain, E12.5 (B) Distribution of H3K36me3 among 6 tissues shows hPTM distribution was significantly different among different types of skipped exons (p-values, ANOVA test).

Comparison of hPTM distribution across different time points and tissues revealed unique patterns for some hPTMs (Supplemental Figures 3-36). When we compared different tissues at the same time point, H3K4me enrichment displayed the biggest variation, e.g. H3K4me3 signal was higher in the exon flanking regions of inclusion gain vs loss category in forebrain at E15.5, while in heart, it was much higher at exons in low vs high category. H3K4me also varied across different time points for some tissues. For example, in heart tissue at E12.5, H3K4me3 enrichment was greatest for exons in the inclusion gain vs loss category, but this preferential enrichment gradually switched to exons in the high vs. low group over developmental time. These results suggested that the role of of hPTMs in AS may vary across time points and tissues.

### Modeling skipped exon inclusion by logistic regression and random forest

To determine if specific histone modifications associated with skipped exon inclusion in different tissues and time points, we took our analysis one step further to computationally model the relationship between histone modifications and each skipped exon category. We tested the hypothesis that the model can distinguish two different groups of skipped exons: (1) exons with developmental gain versus exons with developmental loss and (2) exons with isoform selected high versus exons with isoform selected low. In this study, we chose two different approaches – logistic regression and random forest – because of their good performance and ease of interpretability. For each histone modification, we summed the ChIP-seq signal upstream and downstream (+/- 150bp) of skipped exons’ splice sites and regarded them as 8 hPTM features of the model. These 8 features were then expanded to 32 explanatory variables to build the model. The model performance was measured by accuracy and area under the ROC curve (AUC) based on 5-fold cross validation.

Table 1 shows the accuracy values of two models for 7 tissues at different developmental time points. In general, random forest showed better performance than logistic regression in all tissues and time points, which was consistent with a recent study that compared the performance of 13 popular machine learning algorithms [37]. The accuracy of random forest model varied from 0.57 to 0.72 in developmental gain vs developmental loss category and from 0.67 to 0.70 in isoform selected high vs isoform selected low category. Due to the imbalanced datasets of some tissues, we also compared their AUC values, which is insensitive to imbalanced classes. Consistent with accuracy values, AUC of random forest was between 0.64 and 0.74 in developmental gain vs developmental loss category and from 0.72 to 0.75 in isoform selected high vs isoform selected low category (Figure 3A, Supplement Figures 37-40). In addition, accuracy and AUC values from random forest were much higher than random prediction (0.5), indicating a good predictive power of random forest model.

**Figure 3.**
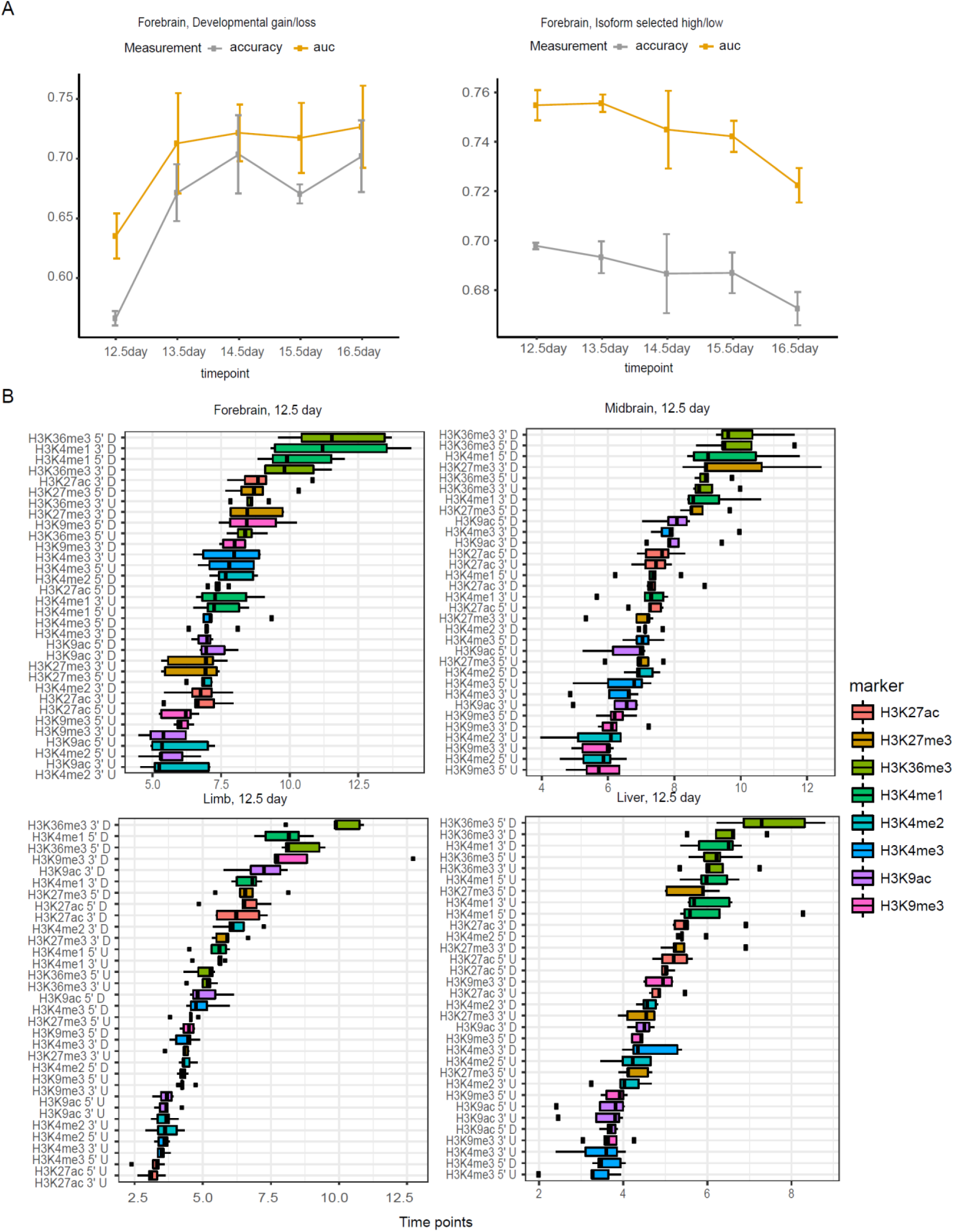
Model performance and important histone modifications associated with skipped exon selection in different tissues and timepoints (A) Accuracy and AUC values of random forest models built based on hPTM signals on the flanking regions of skipped exons in forebrain across developmental time points. (B) H3K36me3 and H3K4me1 are the most predictive hPTMs in differentiating skipped exon inclusion categories. Boxplot of important score generated by random forest model in different tissues at E12.5 shows several types of hPTMs are key predictors. Importance score is calculated based on 5-fold cross validation.

For developmental gain vs loss category, the model performance varied over time. The AUC of random forest was greater than 0.6 for the majority of tissues and time points and showed a trend for increase over time, but this trend may be caused by the smaller sample size at early time points (Figure 3A, Supplemental Figures 37-38). For the skipped exon in isoform selected high vs isoform selected low category, the model performance remained stable for all tissues and did not differ greatly when compared to most of the time points of gain vs loss inclusion category (Table 2, Figure 3A, Supplemental Figures 39-40). In summary, our data indicated an association of hPTMs with two categories of skipped exon selection: exons that show isoform switching behavior during tissue development and exons that are alternatively spliced but without isoform switching over developmental time. In both categories hPTMs were highly predictive of skipped exon inclusion, suggesting that hPTMs are involved in skipped exon selection, either directly or indirectly.

**Table 1.** Accuracies of logistic and random forest models to predict gain vs. loss and high vs. low categories for different tissues at each developmental time point. Accuracies were calculated based on 5-fold cross validation.

| Gain v.s.<br>Loss | Logistic regression |  |  |  |  | Random forest |  |  |  |  |
| --- | --- | --- | --- | --- | --- | --- | --- | --- | --- | --- |
| tissue | 12.5<br>day | 13.5<br>day | 14.5 day | 15.5 day | 16.5 day | 12.5<br>day | 13.5 day | 14.5 day | 15.5 day | 16.5<br>day |
| forebrain | 0.53 | 0.59 | 0.64 | 0.60 | 0.62 | 0.57 | 0.67 | 0.70 | 0.67 | 0.70 |
| heart | 0.62 | 0.62 | 0.50 | 0.62 | 0.60 | 0.62 | 0.64 | 0.64 | 0.64 | 0.62 |
| hindbrain | 0.53 | 0.63 | 0.63 | 0.63 | 0.59 | 0.58 | 0.69 | 0.71 | 0.70 | 0.69 |
| limb | 0.52 | 0.57 | 0.56 | 0.58 | NA | 0.61 | 0.61 | 0.67 | 0.64 | NA |
| liver | 0.59 | 0.54 | 0.47 | 0.54 | 0.53 | 0.64 | 0.59 | 0.52 | 0.59 | 0.61 |
| midbrain | 0.58 | 0.63 | 0.65 | 0.62 | 0.62 | 0.59 | 0.71 | 0.72 | 0.69 | 0.70 |
| Neural tube | 0.55 | 0.61 | 0.65 | 0.59 | NA | 0.59 | 0.67 | 0.72 | 0.68 | NA |

| High v.s. low | Logistic regression |  |  |  |  | Random forest |  |  |  |  |
| --- | --- | --- | --- | --- | --- | --- | --- | --- | --- | --- |
| tissue | 12.5<br>day | 13.5<br>day | 14.5<br>day | 15.5<br>day | 16.5<br>day | 12.5<br>day | 13.5<br>day | 14.5<br>day | 15.5<br>day | 16.5<br>day |
| forebrain | 0.63 | 0.64 | 0.63 | 0.63 | 0.61 | 0.70 | 0.69 | 0.69 | 0.69 | 0.67 |
| heart | 0.61 | 0.63 | 0.61 | 0.62 | 0.62 | 0.69 | 0.70 | 0.69 | 0.70 | 0.68 |
| hindbrain | 0.63 | 0.62 | 0.62 | 0.62 | 0.63 | 0.69 | 0.69 | 0.68 | 0.69 | 0.70 |
| limb | 0.63 | 0.63 | 0.62 | 0.62 | NA | 0.69 | 0.69 | 0.67 | 0.68 | NA |
| liver | 0.62 | 0.60 | 0.60 | 0.59 | 0.59 | 0.69 | 0.67 | 0.67 | 0.66 | 0.67 |
| midbrain | 0.62 | 0.62 | 0.63 | 0.63 | 0.62 | 0.68 | 0.69 | 0.69 | 0.68 | 0.68 |
| Neural tube | 0.65 | 0.63 | 0.62 | 0.64 | NA | 0.70 | 0.69 | 0.67 | 0.70 | NA |

### Specific types of hPTMs were key predictors for skipped exon groups

To elucidate the relative importance of different hPTMs on skipped exon selection and to test if their respective contributions for different tissues and time points, we extracted the importance score generated from random forest model (Figure 3B, Supplemental Figures 41-54). Overall, H3K36me3 and H3K4me1 were the most predictive hPTMs in differentiating skipped exon inclusion categories, while H3K9me3 was the least informative. In addition to H3K36me3, we observed a strong predictive effect of H3K4me1 at 5’ splice site downstream and 3’ splice site upstream in many of the tissues. The 3’ splice site upstream of H3K27me3 showed a greater contribution in midbrain and hindbrain at E12.5, while in limb, H3K9ac and H3K9me3 at the 3’ splice site upstream were informative to differentiate skipped exon groups.

We next compared the same tissue at different developmental time points, and similarly found that H3K36me3 and H3K4me1 ranked at the top for majority of the cases (Supplemental Figures 41-54). Consistent with their contributions at E12.5, the 5’ splice site downstream, 3’ splice site upstream of H3K36me3 and the 5’ splice site downstream, 3’ splice site upstream of H3K4me1 were the most informative predictors. On the other hand, the contribution of some types of hPTMs varied over time. For example, in liver, ChIP-seq signal of H3K9ac at 3’ splice site upstream at E13.5 had a much stronger predictive effect when the same region was compared at the other time points.

To further examine the contribution of individual histone modifications, we averaged the important score in the flanking regions for each hPTM and normalized it by dividing the largest averaged value. We then plotted the normalized score for each time point. Figure 4 shows the contribution of hPTMs to differentiate developmental gain vs. developmental loss category. Similar to what we observed in the previous model, H3K36me3 and H3K4me1 played dominant predictive roles for all tissues over all time points. The contribution of other hPTMs varied across the different tissues examined. In brain tissues and heart, H3K9ac had a relatively higher predictive rank, while in limb, neural tube and liver, the effect of H3K27me3 was greatest. The pattern was consistent for isoform selected high vs. low group (Supplemental Figure 55-56). These results confirmed our hypothesis that specific types of hPTMs associate with skipped exon selection and suggested that H3K36me3 and H3K4me1 play a dominant role in regulating alternative splicing.

**Figure 4.**
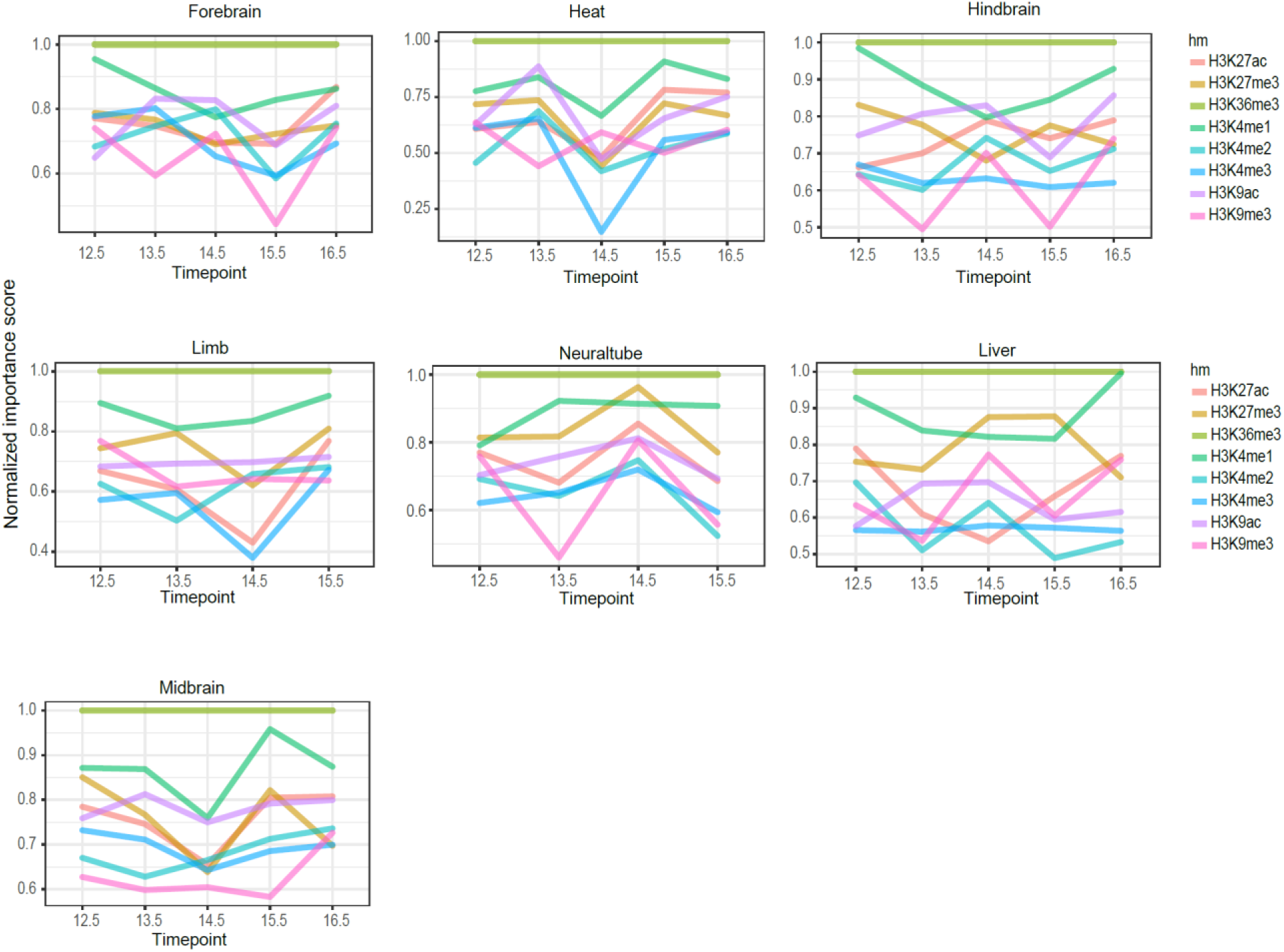
Contributions of different types of hPTMs to differentiate developmental gain versus developmental loss over time in 7 tissues.

### Interaction among histone modifications and skipped exon selection

Interactions among histone modifications in promoter regions for the regulation of gene expression and exon splicing have been reported in several studies based on Bayesian methods [20, 38]. However, Bayesian methods used in these studies discretized the ChIP-seq signal based on the clustering result, which may cause information loss. To investigate the interaction among histone modifications, we used iterative random forest model (iRF), which can be applied to identify high-order interactions [39]. iRF algorithm first sequentially grows feature-weighted random forests to perform feature space reduction and then fits the model based on Random Intersection Trees algorithm to identify high-order feature combinations that are prevalent on the random forest decision paths [39, 40].

As demonstrated in Figure 6, many histone modification interactions were observed in forebrain, heart and liver for developmental gain vs loss category at E12.5. The interactions from other tissues and isoform selected high vs low category are in Supplemental Figures 57-70. These included interactions between modifications on different amino acids (e.g. H3K36me3 and H3K4me1), between different kinds of modifications (e.g. H3K4me1 and H3K9ac), and between the different genic regions of the same histone modification (e.g. H3K4me1 5’ downstream and H3K4me1 3’ upstream). The interaction between H3K36me3 and H3K4me1 in the exon flanking regions (H3K36me3 5’ downstream and H3K4me1 3’ upstream) was the top feature in both developmental gain vs. loss and isoform selected high vs. low group. The other top interactions included H3K27me3 and H3K36me3, H3K27ac and H3K36me3, H3K27ac and H3K4me1 and H3K36me3 and H3K9me3. Interestingly, we observed many interactions between the different flanking regions of the same histone modification, such as interactions between H3K36me3 5’ upstream and H3K36me3 3’ upstream, suggesting a spatial relevance of hPTMs to alternatively spliced exon selection.

**Figure 5.**
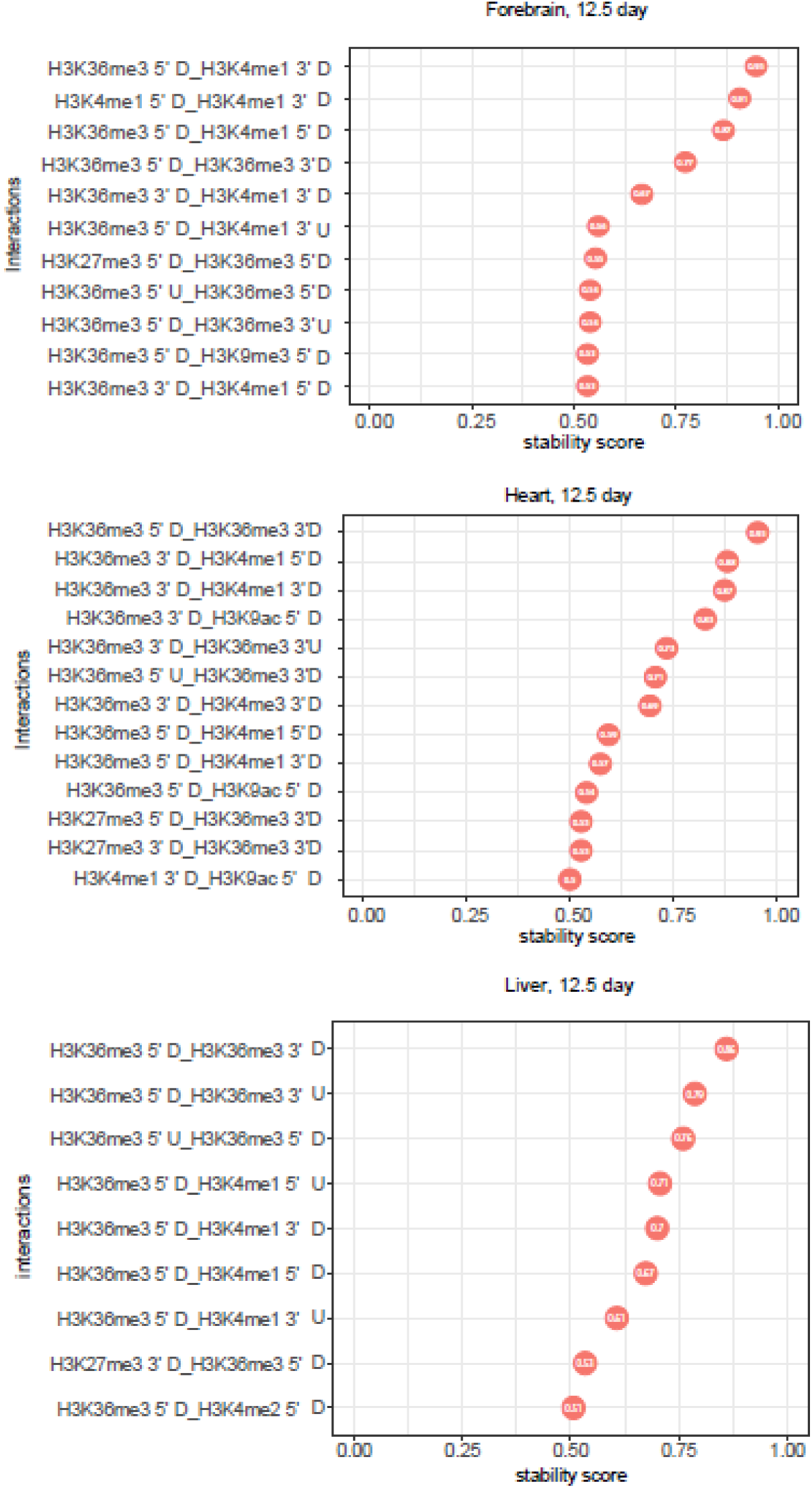
Top stable interactions of histone modifications in the exon flanking region for gain vs loss group obtained by iRF algorithm (stability score >= 0.5). 3 tissues (12.5 day) were visualized: forebrain, heart and liver.

## Discussion

AS plays a critical role during tissue development and cell differentiation [2, 29]. Previous studies reveal several regulatory mechanisms for AS, including expression and targeting of splicing factors and enrichment of hPTMs [29]. In this study, we sought to comprehensively investigate previous observations of the relationship between hPTMs and AS during tissue development by integrating ChIP-seq and RNA-seq data from 7 different mouse embryonic tissues at 6 developmental time points. We identified two different categories of AS (skipped) exons: skipped exons associated with development and skipped exons associated with isoform selection. Ontological analyses found that genes from these two categories are enriched in different functions. Developmentally associated AS genes in forebrain were more likely to be enriched in neuronal-related functional categories, such as neuron projection, postsynaptic density and cytoskeleton. This is consistent with previous gene ontology analysis for differentially spliced exons in developing cerebral cortex, which show that cytoskeleton genes are overrepresented in mouse and human [34]. On the other hand, AS genes associated with isoform selection are overrepresented in distinct ontological categories important to maintain cell function or tissue identity, such as cell cycle, protein transport and RNA binding.

Computational models constructed based on ChIP-seq signal in the flanking regions of skipped exons showed that hPTMs associated with both categories of skipped exon. Consistent with previous studies, we found H3K36me3 to be most predictive for skipped exon groups [36, 41, 42]. Specifically, H3K36me3 enrichment in the flanking region 5’ splice site downstream and 3’ splice site upstream of the skipped exon was the top predictor in all tissues, which implied the dominant role of H3K36me3 in skipped exon selection. This result has been reported previously in other systems. First, a computational analysis of skipped exons based on 3 human cell lines shows the enrichment of H3K36me3 in exon and downstream of 3’ splice site is significantly correlated with skipped exon inclusion [36]. Second, analysis of human and *C. elegans* exons finds H3K36me3 is most strongly enriched at a well-positioned nucleosome located at the 5’ end of exons [43]. The stronger enrichment of H3K36me3 correlates with increased exon usage in alternatively spliced genes. Third, a recent study that analyzed transcriptomic data from human embryonic stem cells at various stages of differentiation shows that H3K36me3 is the top important feature to identify splicing codes across different cell lineages and datasets [27]. Finally, our previous analysis of mouse nucleus accumbens reveals that H3K36me3 has the greatest enrichment at alternative isoforms relative to other hPTMs [44]. In addition to these computational approaches, causal molecular mechanisms for the role of H3K36me3 have also been elucidated in the literature. Luco et al validates the causal relationship between hPTM and alternative splice site selection using exogenous enrichment of H3K36me3 and co-immunprecipiation experiments in which H3K36me3 is shown to physically interact with PTB protein to regulate alternative splicing [45]. Together with our findings, these prior reports indicate one possible regulatory by which H3K36me3 enrichment contributes to alternative exon selection in AS (Figure 6).

**Figure 6.**
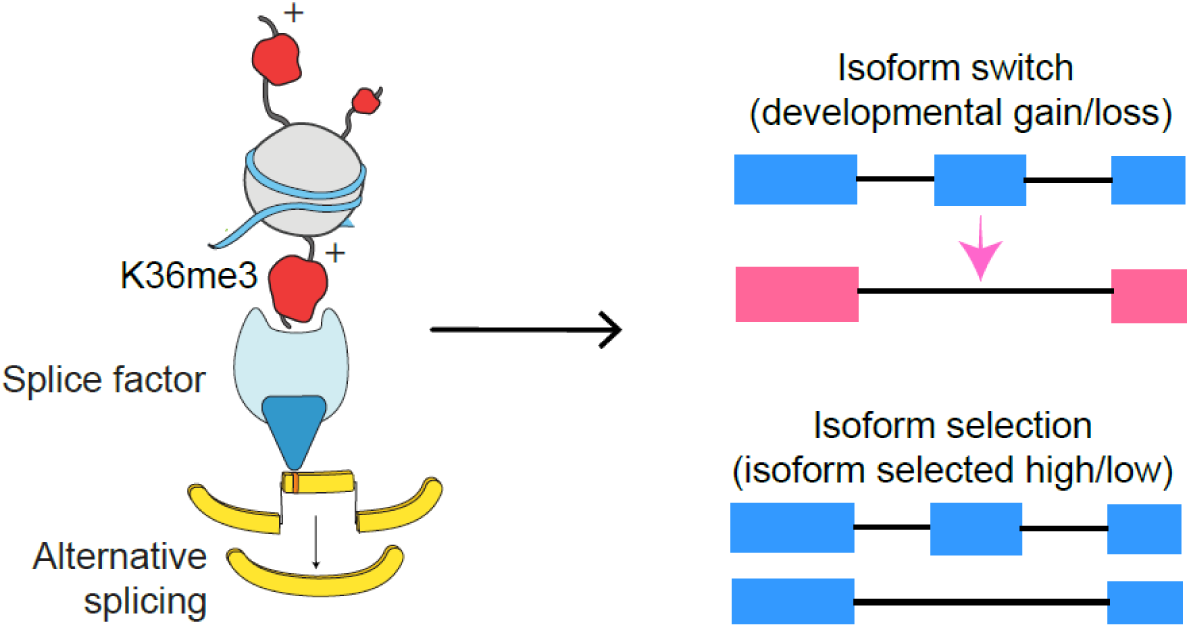
Regulatory model indicating how a particular hPTM (e.g. H3K36me3) regulates skipped exon selection during alternative splicing.

Using iterative random forest model, we identified interactions between several hPTMs that associated with skipped exon selection, including interactions between modifications on different amino acids (e.g. H3K36me3 and H3K4me1), between different kinds of modifications (e.g. H3K4me1 and H3K9ac), and between the different genic regions of the same histone modification. The interaction between H3K36me3 and H3K4me1 in the exon flanking regions was the top feature in both developmental gain vs. loss and isoform selected high vs. low group. Other interactions, such as H3K27me3 and H3K36me3, H3K27ac and H3K36me3, and H3K36me3 and H3K9me3, indicated that relatively weaker predictive hPTMs may only be functional when in combination with the highly predictive ones. The concept of hPTM interaction is not new. Several studies have found the combinatorial effect of histone modifications and their association with gene transcription and differentiation. For example, Jaschek et al measure the co-occurrence of many histone modifications in the region of transcription start site [46]. Another similar study by Han et al deciphers histone modification interaction relationships on exons based on Bayesian network [20]. Interestingly, we found many interactions that occur between the different flanking regions of same or different histone modifications, such as interactions between H3K36me3 5’ upstream and H3K36me3 3’ upstream and between H3K36me3 5’ downstream and H3K4me1 3’ upstream. These results suggested hPTMs located in different positions in the exon flanking region may contribute differently to skipped exon selection. This is consistent with the result of a previous study that finds hPTMs correlate to skipped exon inclusion via specific patterns along the flanking region of those exons [22]. In particular, H3K36me3 shows a significant correlation between upstream and downstream of exon flanking regions for exon inclusion rate, which is consistent with our finding.

Furthermore, we observed the occurrence of some hPTMs at several skipped exons, suggesting a potential mechanistic connection between those modifications and tissue development driven by AS (Supplemental table 4). Many of these multi-hit exons, such as Filamin-A protein (*FLNA*), were also identified in a separate study of the developing cerebral cortex [34]. RNA-seq analysis from Zhang et al shows *FLNA* exonN is included in cerebral cortex and cerebellum but excluded from non-neural tissues, which is consistent with our finding that the inclusion level of *FLNA* exonN is significantly increased in forebrain over developmental time, but not in liver, heart and limb. Interestingly, one previous study finds that mutations that disrupt the Polypyrimidine tract binding protein (PTBP1) binding site of *FLNA* exonN in neural progenitor cell causes a brain-specific malformation in human, suggesting the potential regulatory role between PTBP1 and exonN inclusion [34]. In this study, we observed that the signal of H3K36me3 in the flanking regions was significantly correlated with *FLNA* exonN inclusion in forebrain (Figure 7A, B), which may indicate the potential link among histone modification, splicing factor and exonN inclusion (Figure 7C). This notion is furthered by the previous findings from Luco et al, that H3K36me3 can directly interact with spliceosome components, PTB and MRG15, regulating the expression of target exons at FGFR2, TPM2, TPM1, and PKM2 loci in human cell lines [45].

Overall, we have performed a comprehensive analysis to investigate chromatin-mediated alternative splicing events during tissue development. Using computational models, we found that specific histone modifications, H3K36me3 and H3K4me1, played a dominant role in skipped exon selection among all the tissues and developmental time points examined. We also identified interactions of two or more hPTMs that highly predict AS. For example, the interaction between H3K36me3 and H3K4me1 in the exon flanking region was the top feature in both skipped exon categories. These findings increased the complexity of defining AS regulation. Our data demonstrated a link between hPTMs and alternative splicing in mouse tissue development, which will drive further experimental studies on the functional relevance of these modifications to alternative splicing.

**Figure 7.**
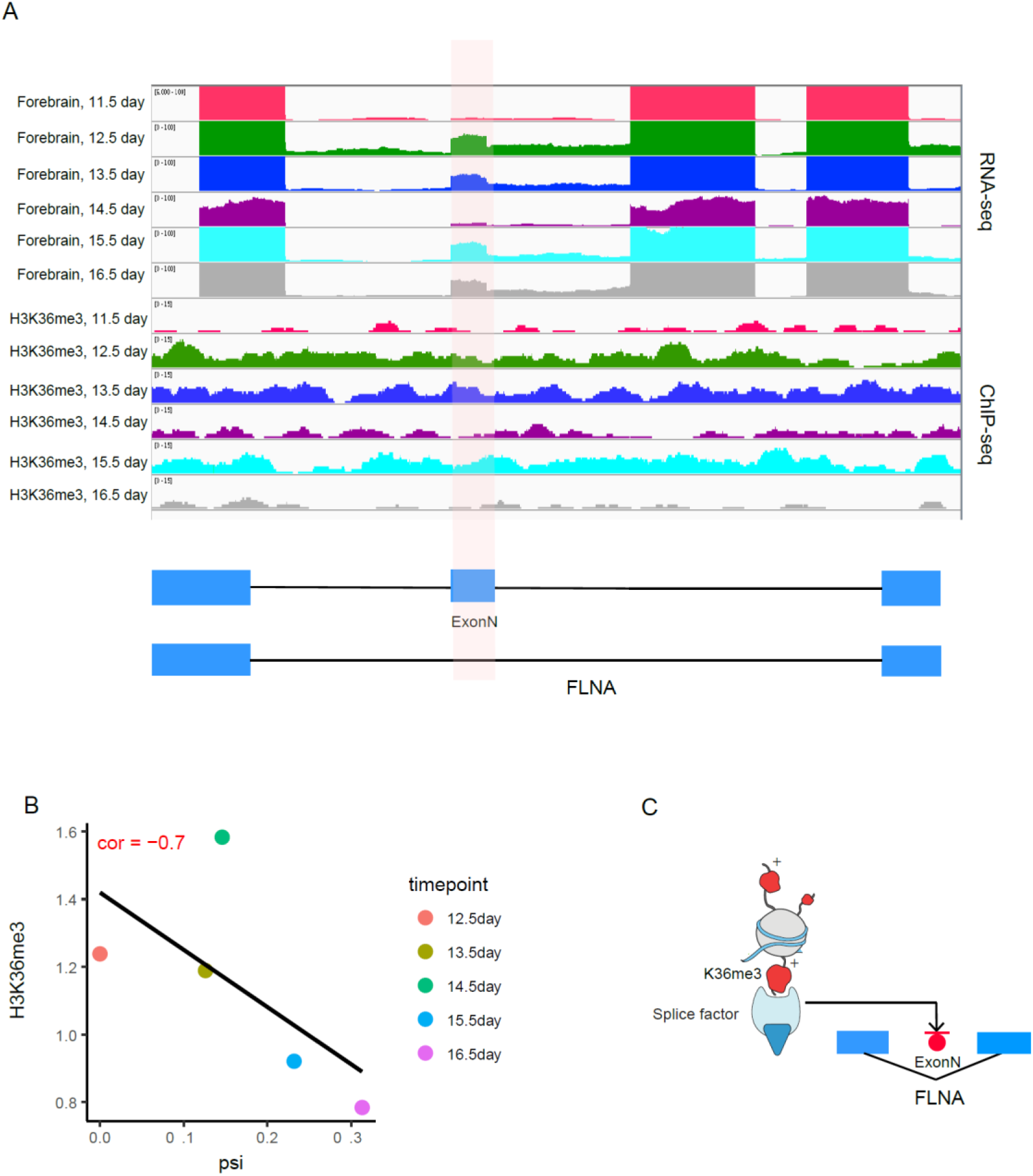
Potential mechanistic connection between histone modifications and tissue development. (A) Genome browser view shows the enrichment of H3K36me3 and exonN inclusion in forebrain, heart, liver and limb. (B) The inclusion level of ExonN in *FLNA* gene was significantly correlated with H3K36me3 enrichment. (C) Schematic depicts a potential mechanism by which H3K36me3 regulates exonN inclusion. H3K36me3 can recruit splicing factor PTBP1, which will further repress exonN inclusion.

## Methods

### Dataset

We chose mouse embryonic tissue developmental data from ENCODE database [30], because both RNA-seq and ChIP-seq are available. We considered 7 tissues (forebrain, hindbrain, midbrain, neural tube, heart, liver and limb) from 6 time points (E11.5 – E16.5 day). Supplemental Table 1 provides the full list of data analyzed. The analysis codes are available through github https://github.com/huqiwen0313/HM_splicing.

### Identification of alternative splicing exons in tissue development

Aligned BAM files (mm10) for all 7 tissues from 6 timepoints were downloaded from ENCODE 455 [30], each with two replicates. rMATS (version 4.0.1) was used to quantify ‘percent spliced in’ (PSI, exon inclusion level) and identify skipped exons that showed differential inclusion level (deltaPSI) between two time points [31]. Skipped exons were divided into two different groups based on PSI and ΔPSI values: gain versus loss and high versus low. For gain versus loss group, we chose exons with ΔPSI >= 0.01, p-value <= 0.01 and FDR <= 5% as gain class and exons with ΔPSI <= 0.01, p-value <= 0.01 and FDR <= 5% as loss class. For high versus low group, we generated the global PSI distribution for all skipped exons. The upper (75%) and lower (25%) quantiles are used as the cutoffs to divide exons into high class and low class. Exon inclusion level change between two different time points for one tissue identified skipped exons in gain versus loss group. These exons showed the isoform switch behavior during tissue development. On the other hand, skipped exons in high versus low group refer to those exons that were alternative spliced, but their exon inclusion levels did not change over time.

### ChIP-seq data processing and hPTM profiling

ChIP-seq data (aligned BAM files, mm10) were downloaded from ENCODE database [30]. For each tissue and time point, 8 type of histone modifications, including H3K36me3, H3K4me1, H3K4me2, H3K4me3, H3K27ac, H3K27me3, H3K9ac and H3K9me3 were analyzed. The global profiles of histone modification among different groups of skipped exons were generated in two steps. First, for each exon, the flanking regions were defined as the 300bp centered at acceptor and donor site, respectively, analyzed in 15bp bins. Second, the ChIP-seq reads were assigned to those binned regions and the normalized reads number for each binned region was calculated. To visualize the ChIP-seq signal pattern for different exon groups, we computed the average ChIP-seq signal across the flanking regions, averaged over all exons that belong to the same group. An ANOVA statistic was used to test if the signal distribution patterns were significantly different among different skipped exon groups.

### Logistic regression and random forest modelling

To extract the features from hPTM distribution patterns, the flanking regions surrounding each splice site were divided into 4 regions: the intronic region at the acceptor splice site (5’ upstream, left_intron), the exonic region at the acceptor splice site (5’ downstream, left_exon), the exonic region at the donor splice site (3’ upstream, right_exon) and the intronic region at the acceptor splice site (5’ downstream, right_intron). The normalized ChIP-seq signals in those regions were calculated and considered as explanatory features for different types of hPTMs.

To demonstrate a predictive association between ChIP-seq signal and skipped exon groups, we constructed binary classification models. We chose two different models: logistic regression and random forest. Logistic regression is a type of probabilistic statistical classification model that measures the relationship between categorical response variable and explanatory variables, which can be formulated as below:

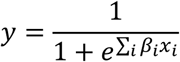

in which x_i_ is the ChIP-seq signal for certain type of hPTM, y is skipped exon groups and β_i_ is the regression coefficient.

Random forest is an ensemble tree-based algorithm that uses bootstrap resampling to grow multiple decision trees and combines their results. The advantage of logistic regression and random forest over the other models is the interpretability of the model results, that is, we can know the effect of individual feature to the response variable.

The model performance was measured by 5-fold cross validation, in which the entire datasets was randomly partitioned into 5 equal-sized subsamples. One subsample was used to evaluate the model performance (test set) and the remaining subsamples (training set) were used to train the model. The whole process was repeated by 5 times. Average model accuracy and ROC value were then calculated.

To generate statistical robustness, for each training set, the model was further tuned by a grid of parameters based on internal 3-fold cross validation. The model with the lowest error rate was then selected. For logistic regression, in order to achieve better performance, LASSO was applied to reduce the dimension of feature space, because when the feature space is large, the ordinary least square estimates generated by logistic regression may lead to large variance for the estimates, which will reduce the accuracy of prediction. We estimated the LASSO parameter λ through 3-fold cross validation. For each cross validation, a grid of λs was fed to the model. The corresponding prediction was estimated according to the test set. The λ value that minimized the overall prediction error was selected.

### Iterative random forest modelling and interaction analysis

Iterative random forest model searched for high-order interaction in three steps: 1. Iteratively re-weighted random forests. 2. Extract decision rules from feature-weighted random forest path and recover interactions. 3. A bagging step to assess the stability of interactions. The detailed description of iterative random forest algorithm is in [39]. We trained iterative random forest model using R package iRF (https://github.com/sumbose/iRF), with number of iterations = 10 and number of bootstraps = 30. The stability score was estimated through 5-fold cross validation. Interactions with stability score > 0.5 were considered as meaningful interactions.

### Gene ontology enrichment analysis

The Gene Ontology (GO) enrichment analysis was performed using DAVID [47] under default parameters. Overrepresented GO terms for GO domain belong to biological process, cellular component and molecular function were used to generate enrichment dataset based on FDR cutoff 0.05.

